# Insecticide resistance and its intensity in urban *Anopheles arabiensis* in Kisumu City, Western Kenya: *Implications for malaria control in urban areas*

**DOI:** 10.1101/2024.05.06.592663

**Authors:** Maxwell G. Machani, Irene Nzioki, Shirley A. Onyango, Brenda Onyango, John Githure, Harrysone Atieli, Chloe Wang, Ming-Chieh Lee, Andrew K. Githeko, Yaw A. Afrane, Eric Ochomo, Guiyun Yan

**Affiliations:** Centre for Global Health Research, Kenya Medical Research Institute, Kisumu, Kenya; School of Zoological Sciences, Kenyatta University, Kenya; International Center of Excellence for Malaria Research, Tom Mboya University, Homa Bay, Kenya; Department of Medical Microbiology, University of Ghana Medical School, College of Health Sciences, University of Ghana, Ghana; Program in Public Health, College of Health Sciences, University of California, Irvine, CA 92697, USA

**Keywords:** *Anopheles*, insecticide resistance intensity, pesticide use, urban, western Kenya

## Abstract

**Background:** The rise of insecticide resistance poses a growing challenge to the effectiveness of vector control tools, particularly in rural areas. However, the urban setting has received comparatively less focus despite its significance in attracting rural to urban migration. Unplanned urbanization, often overlooked, exacerbates insecticide resistance as *Anopheles* mosquitoes adapt to the polluted environments of rapidly expanding cities. This study aimed to assess the insecticide susceptibility status of malaria vectors and identify potential underlying mechanisms across three distinct ecological settings characterized by differing levels of urbanization in Kisumu County, Kenya.

**Methods:** Field-derived *An. gambiae* (s.l.) larvae collected from a long stretch of urban-to-rural continuum were phenotyped as either resistant or susceptible to six different insecticides using the World Health Organization (WHO) susceptibility test. Polymerase chain reaction (PCR) techniques were used to identify the species of the *An. gambiae* complex and screened for mutations at voltage-gated sodium channels (Vgsc-1014F, Vgsc-1014S, Vgsc-1575Y) and acetylcholinesterase Ace1-119S. Metabolic enzymes activities (non-specific β-esterases and monooxygenases) were evaluated in mosquitoes not exposed to insecticides using microplate assays. Additionally, during larval sampling, a retrospective questionnaire survey was conducted to determine pesticide usage by the local inhabitants.

**Results:** *Anopheles arabiensis* dominated in urban (96.2%) and peri-urban (96.8%) areas, while *An. gambiae* (s.s.) was abundant in rural settings (82.7%). Urban mosquito populations showed high resistance intensity to deltamethrin (Mortality rate: 85.2% at 10x) and suspected resistance to Pirimiphos-methyl and bendiocarb while peri-urban and rural populations exhibited moderate resistance intensity to deltamethrin (mortality rate >98% at 10x). Preexposure of mosquitoes to a synergist piperonyl butoxide (PBO) significantly increased mortality rates: from 40.7% to 88.5% in urban, 51.9% to 90.3% in peri-urban, and 55.4% to 87.6% in rural populations for deltamethrin, and from 41.4% to 78.8% in urban, 43.7% to 90.7% in peri-urban, and 35% to 84.2% in rural populations for permethrin. In contrast, 100% mortality to chlorfenapyr and clothianidin was observed in all the populations tested. The prevalence of L1014F mutation was notably higher in urban *An. arabiensis* (0.22) unlike the peri-urban (0.11) and rural (0.14) populations while the L1014S mutation was more prevalent in rural *An. gambiae* (0.93). Additionally, urban *An. arabiensis* exhibited elevated levels of mixed function oxidases (0.8/mg protein) and non-specific esterases (2.12/mg protein) compared to peri-urban (0.57/mg protein and 1.5/mg protein, respectively) and rural populations (0.6/mg protein and 1.8/mg protein, respectively). Pyrethroids, apart from their use in public health through LLINs, were being highly used for agricultural purposes across all ecological settings (urban 38%, peri-urban 36% and rural 37%) followed by amidine group, with organophosphates, neonicotinoids and carbamates being of secondary importance.

**Conclusion:** These findings show high resistance of *An. arabiensis* to insecticides commonly used for vector control, linked with increased levels of detoxification enzymes. The observed intensity of resistance underscores the pressing issue of insecticide resistance in urban areas, potentially compromising the effectiveness of vector control measures, especially pyrethroid-treated LLINs. Given the species’ unique behavior and ecology compared to *An. gambiae*, tailored vector control strategies are needed to address this concern in urban settings.

## Background

The implementation of insecticide-based control strategies, such as long-lasting insecticide-treated nets (LLINs) and indoor residual spraying (IRS), has notably reduced malaria-related morbidity and mortalities in sub-Saharan Africa (1). However, the survival of malaria vectors despite these interventions has become a growing challenge due to increasing insecticide resistance and behavioral avoidance (2). This challenge is further exacerbated by urban malaria, which has emerged as a significant concern due to ongoing rural-to-urban migration, and rapid unregulated urban development, allowing malaria vectors to adapt to a wide range of polluted habitats (3). While insecticide resistance poses an emerging threat to the efficacy of LLINs and IRS in rural areas, relatively less attention has been directed towards addressing this issue in urban settings. Urban environments are generally known to have lower populations of *Anopheles* mosquitoes compared to rural areas, primarily due to elevated levels of human pollution and disturbance (4). Due to increasing human population pressure and urbanization witnessed in recent decades, significant ecological changes have unfolded in most African cities, influencing the distribution of vector species (5, 6). Unplanned urbanization often results in a proliferation of man-made breeding sites, many of which contain higher levels of chemical pollutants or other xenobiotics compared to rural areas (7, 8). *Anopheles gambiae* (s.l.), which traditionally preferred unpolluted water sources (9), has now exhibited a notable adaptation to polluted waters in urban areas (8, 10). Furthermore, these mosquitoes are now breeding in diverse human-made habitats, including containers (11). The presence of anthropogenic pollutants in water bodies (12), may indirectly contribute to the selection of mosquito resistance to chemical insecticides. While these urban pollutants might not be directly toxic to mosquitoes, they can swiftly influence the mosquito’s resistance to different insecticides, primarily by inducing detoxification enzyme activities (13). In laboratory settings, mosquito larvae exposed to anthropogenic chemicals have demonstrated an enhanced ability to survive insecticide exposure (14). Understanding the resistance status of vectors in urban areas against insecticides used for vector control and the underlying mechanisms is crucial when devising an integrated vector control strategy for effective vector management in urban settings.

Pyrethroids, until recently were the only class of insecticides recommended for use on LLINs. However, pyrethroid resistance amongst *Anopheles* mosquitoes has become widespread (15). Two main resistance mechanisms, conferred by genomic mutations, are now widely documented across Africa: (1) target site modifications and (2) enhanced detoxification of insecticides (16). Additionally, other resistance mechanisms, such as cuticular thickening and overexpression of chemosensory proteins, have been reported (17, 18). Moreover, there is evidence of malaria vectors exhibiting key behavioral shifts that render them less susceptible to LLINs (19–21). The increase in resistance has been linked to enormous selection pressures resulting from large-scale and regular deployment of these insecticides (22–24). Besides, the domestic application of insecticides at the household level and the use of the same classes of insecticides for agricultural purposes have been documented to influence local variation in insecticide resistance (25). In Kenya, over the past decade, malaria vector resistance to the four classes (pyrethroids, organochlorines, carbamates and organophosphates) of insecticides has intensified (26, 27). Metabolic resistance mediated by mixed-function oxidases and the presence of the kdr mutation are prevalent mechanisms associated with insecticide resistance. The L1014S mutation is fixed, indicating it is widespread occurrence, while the L1014F mutation has been identified but at considerably low frequencies (28, 29).

It’s noteworthy that previous studies in western Kenya have primarily focused on assessing resistance levels and associated mechanisms in rural areas where malaria is endemic (26–28, 30, 31). However, the impact of the observed adaptation on mosquito tolerance to commonly used insecticides in vector control and the impact of urban agricultural within the urban environment remains unknown. While controlling urban malaria vectors may involve targeting the immature stages through measures such as larviciding and environmental management (32), the primary control measure in Kenya typically revolves around preventing transmission using LLINs. Therefore, this study was conducted to evaluate the resistance status of malaria vectors to insecticides used in vector control across different ecological profiles (urban, peri-urban and rural) and to identify potential underlying mechanisms.

## Methods

### Study sites

The study was conducted across three ecological zones in Kisumu County: urban, peri-urban, and rural areas. The county’s urbanization revolves around the city of Kisumu. Urban Kisumu (00°06′S 034°45′E) lies between east of Lake Victoria with an elevation of approximately 1,140m above sea level and west of the Nyando Escarpment. Thirteen sites were randomly selected along an urban-to-rural transect from Kisumu city spanning a distance of 30km. Among these, five locations were surveyed within the city and characterized by dense urbanization (Nyalenda, Gesoko, Migosi, Mamboleo and Bandani, all are informal residences located within the Kisumu city), while four sites were peri-urban (Kotetni, Kandalo, Tiengre and Kisian) and another four were rural (Ojola, Mainga, Chulaimbo and Marera, approximately 30 km from the city) (Fig 1). The zones were classified according to their levels of development and the system of physical planning. Kisumu experiences a humid climate with an average relative humidity of 70% and two distinct seasons: a long rainy season from March to May and short rains in September to December. The extended dry season spans from January to March, with a shorter dry period from August to September. Annual rainfall typically ranges between 1,000 and 1,500 mm. *Anopheles* mosquito species in the region include *Anopheles arabiensis*, *An. funestus*, and *An. gambiae* (27, 33). The region is considered to have low insecticide resistance levels (28, 34).

**Figure 1.**
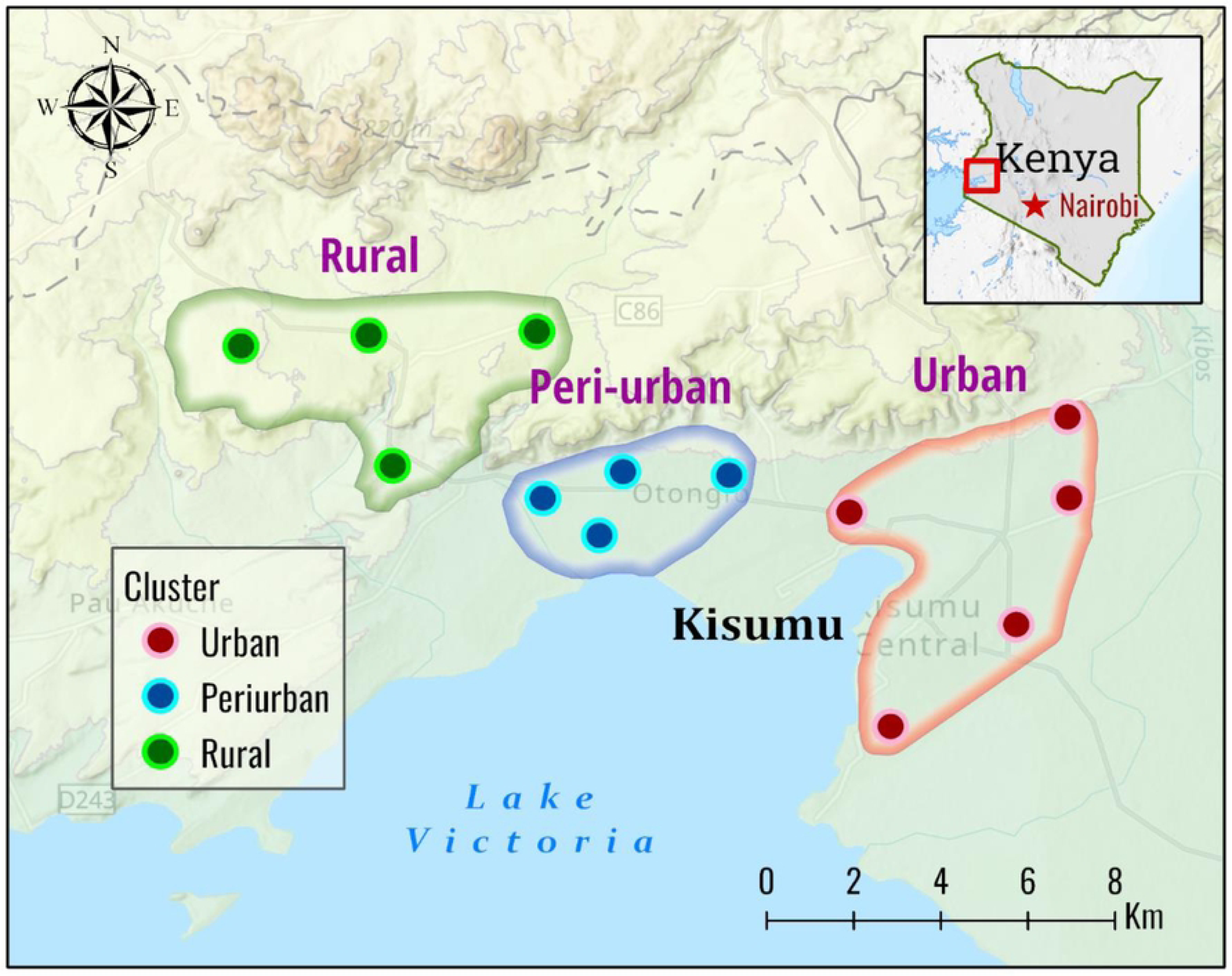
Map of Kenya (right corner) and Kisumu County (in expanded view) showing mosquito collection sites (circles) in the three zones (urban, peri-urban and rural areas in western Kenya (The map was generated using ArcGIS Pro 2.6 software. Map source: ESRI, CGIAR, and USGS (available at: www.esri.com).

### Anopheline mosquito sampling

Anopheline mosquito larvae were collected from several field surveys, allowed to grow in the insectary to become adults, and tested between August 2022 to December 2023. Sampling was conducted within individual sites measuring 1.5×1.5 km^2^. To ensure genetic diversity and prevent sampling bias from a single mosquito egg deposition, larvae were collected from various breeding habitats in each site (Fig 2). The aquatic stages were pooled per site in each zone and transported to the insectary at the Centre for Global Health Research, Kenya Medical Research Institute (KEMRI) in Kisumu, Western Kenya. Larvae were reared under controlled conditions (25 ± 2 °C; 80% ± 4% Relative Humidity with a 12 h: 12 h light/dark cycle). The larvae were raised in rainwater-filled small trays and provided with daily access to powdered Tetramin® fish food and brewer’s yeast. Upon pupation, individuals were collected and transferred to cages, allowing them to emerge as adults. The emerging adults were provided with a 10% sugar solution until they were ready for use in bioassay tests. For the bioassay tests, the *Anopheles arabiensis* Dongola strain susceptible mosquito colony served as a control. During the larval sampling process, a retrospective questionnaire survey was carried out among residents and landowners (63 urban, 80 peri-urban, and 84 rural households) to assess pesticide usage in farms.

**Figure 2.**
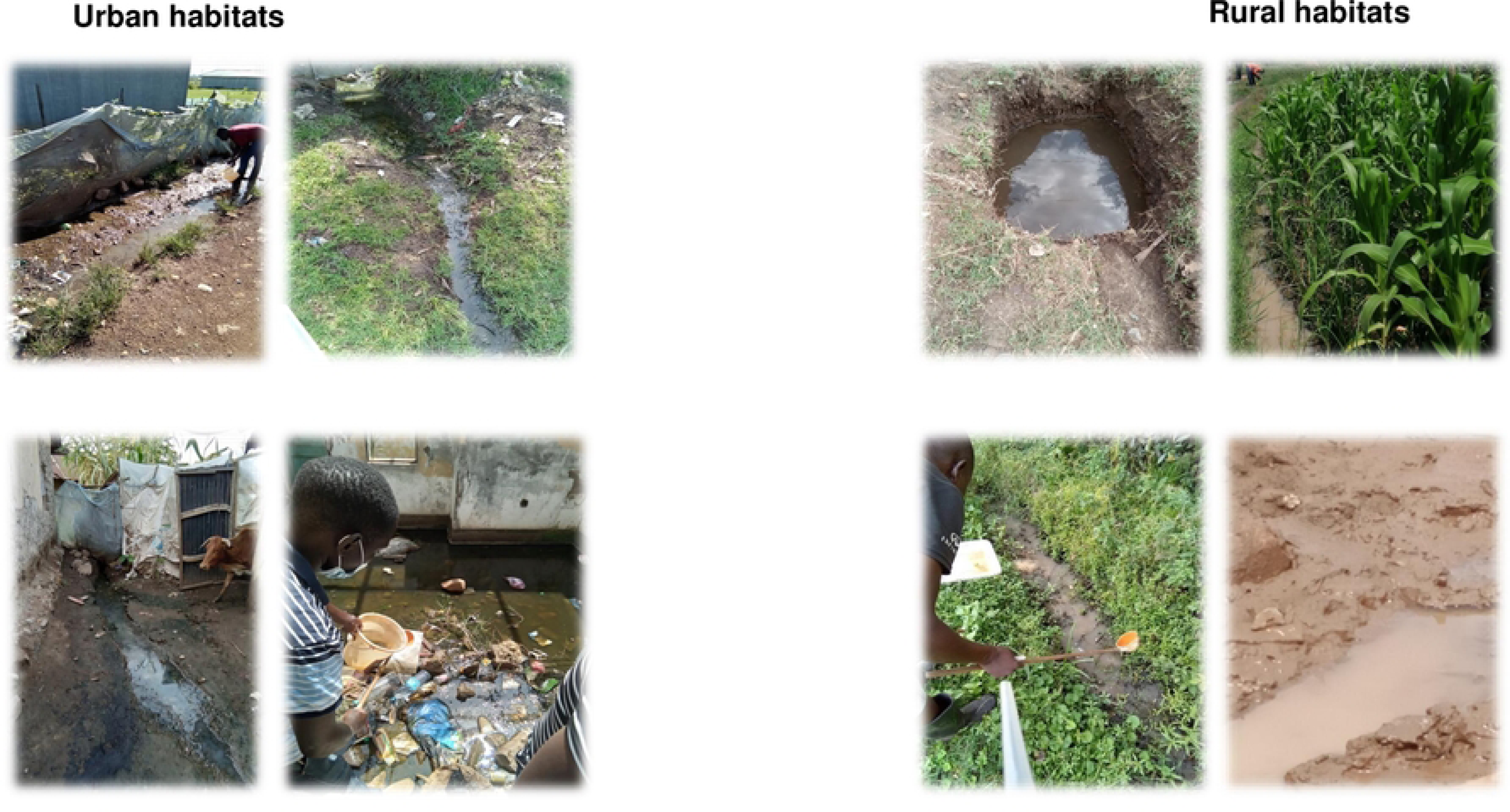
Examples of the habitat types where *Anopheles gambiae* s.l. were collected in urban (left panel) and rural (right panel) zones.

### WHO insecticide susceptibility testing

To assess the insecticide susceptibility of *An. gambiae* (s.l.) populations across the three ecological zones, adult mosquitoes underwent testing using WHO-approved kits following the standard procedures(35). All mosquitoes used in the bioassay were reared from larvae, ensuring uniform age, nutritional status and known physiological status (non-blood-fed). The tests were performed on batches of 25 female anopheline mosquitoes aged 3 to 4 days. The WHO-treated test papers included pyrethroids (0.75% permethrin and 0.05% deltamethrin), organophosphates (0.25% Pirimiphos-methyl), and carbamates (0.1% bendiocarb). Mosquitoes were exposed to the treated papers for 60 minutes, with knockdown rates recorded every 10 minutes. Following the exposure, mosquitoes were transferred to holding tubes and maintained on 10% sugar solutions for 24 hours under controlled conditions (Temperature: 25 ± 2 °C; Relative humidity: 80% ± 4%). Mortality rates were recorded 24 hours post-exposure to the insecticide, and resistance status was determined according to the WHO guidelines on insecticide susceptibility (36). The Sudanese susceptible *An. arabiensis* Dongola strain served as the primary test control to validate the quality of impregnated papers.

### Resistance intensity bioassay

The assessment of resistance intensity for pyrethroids involved using concentrations of 5× and 10× higher than the standard discriminating concentration for deltamethrin (0.05%). Only mosquito populations confirmed to be resistant were subjected to the evaluation of resistance intensity. The test was identical to the standard WHO tube test except that 5× and 10× concentrations were used in the test papers. Four replicates, each consisting of 25 female adult mosquitoes aged 3–4 days, were used for the bioassays. Testing procedures and the determination of resistance intensity categories (i.e., low, moderate and high intensity) followed the standard WHO protocol (35).

### Synergy bioassays

To investigate the role of metabolic detoxification in pyrethroid resistance, piperonyl butoxide (PBO), a synergist that inhibits the specific activity of P450 monooxygenases in insects, was incorporated into the resistance bioassay. This was conducted upon confirming mosquitoes’ resistance to permethrin and/or deltamethrin. Briefly, unfed females aged 3–5 days underwent pre-exposure to 4% PBO-impregnated test papers for 1 hour, followed immediately by exposure to either 0.05% deltamethrin or 0.75% permethrin for an additional hour. A control group of 25 females was only exposed to 4% PBO without any of the insecticides. Subsequently, mosquitoes were transferred to holding tubes and provided with a 10% sugar solution. Mortality rates were recorded after the 24-hour recovery period.

### Determination of susceptibility to chlorfenapyr and clothianidin

Following WHO guidelines (35), bottle bioassays were carried out using four replicates, each consisting of 25 female mosquitoes aged 3–5 days. These mosquitoes were exposed to chlorfenapyr (100 µg/ml, solvent acetone) and clothianidin (4 µg/ml, solvent acetone with MERO) for 1 hour, approximately 24 hours after coating the bottles with the respective insecticides. An additional single replicate of 25 mosquitoes served as a negative control, with bottles treated with 1 ml acetone for chlorfenapyr or acetone with MERO for clothianidin. The knockdown rate was recorded every 10 minutes during the 60-minute exposure period. After exposure, mosquitoes were transferred to a paper cup covered with untreated netting, provided with lightly moistened cotton wool containing a 10% sugar solution (changed daily), and monitored at 24 hours, 48 hours, and 72 hours. Mortality rates were recorded at these time points. Tests were conducted in the morning, with efforts made to maintain testing and holding conditions within WHO guidelines of 27 °C ± 2 °C and a relative humidity of 75% ± 10% (36). The Dongola susceptible strain of *An. Arabiensis* was utilized as the primary test control to ensure the quality of coated bottles.

### Molecular identification of members of the *Anopheles gambiae* complex

Genomic DNA was extracted from individual mosquitoes following previous published methods (37). Molecular identification of sibling species of respective *An. gambiae* (s.l.) were conducted based on PCR methods described by Scott et al. (38).

### Detection of knockdown resistance (*kdr*) and *Ace-1* alleles

Mutations associated with knockdown resistance (*kdr*), L1014S and L1014F, in *An. gambiae* (s.l.) to pyrethroids were assayed using standard TaqMan^®^ quantitative real-time polymerase chain reaction technique (qPCR) (39), using aliquots from the DNA samples extracted for species identification. The same sample sets were genotyped for G119S mutation in *Ace*-1(40).

### Metabolic enzyme activity assays

Adult female mosquitoes emerging from field-collected larvae (aged 3–5 days) were analyzed for activity levels of non-specific β-esterases and mixed function oxidases enzymes. Samples were collected from each study locality, and they had not been exposed to any insecticide prior to the assays. Following Bradfold method (41) for each enzyme, all assays were conducted in triplicate, alongside the Dongola strain serving as a susceptible control population. Briefly, individual whole adult mosquitoes were homogenized in potassium phosphate (KPO_4_) buffer, and substrates specific to each enzyme, along with chromogenic agents, were added. Absorbance values were measured using a microplate reader at wavelengths specific to the enzyme being measured: 540 nm for β-esterases in the presence of β-naphthyl acetate and 620 nm for monooxygenase (cytochrome P450) using 3,3′,5,5′-Tetramethyl-Benzidine Dihydrochloride (TMBZ). Total protein from individual mosquitoes was analyzed to standardize the mean enzyme activity of the test samples.

### Data analysis

Mosquitoes were considered susceptible if mortality was ≥98% and resistant if mortality was <90%. Mortality rates between 90% and 97% indicated possible resistance, requiring confirmation through additional tests (WHO 2016). Resistance intensity was categorized into three (low, moderate and high) following WHO criteria (35). Briefly, mortality < 90% after exposure to 1× DC and ≥ 98% after exposure to 5× DC indicates low-intensity resistance. Mosquitoes with < 90% mortality after exposure to 1× DC, < 98% after exposure to 5× DC, and ≥ 98% at 10× DC indicates moderate intensity. Abbott’s formula was used to correct mortality rates when 5–20% of individuals died in the corresponding control tests (36). Mortality < 90% after exposure to the 1× DC and < 98% after exposure both to 5× and 10× DC indicates high intensity. The allele frequencies for resistant genotypes were calculated using the Hardy-Weinberg equilibrium equation. Mosquito populations were checked to determine whether they were in Hardy-Weinberg equilibrium using *χ* ^2^ test. Biochemical assay data activities (enzymatic activity per mg of protein) were compared to the Dangola *An. arabiensis* reference strain by the Kruskal-Wallis test with Dunn’s multiple comparisons post hoc. In all analyses, *P* < 0.05 was considered significant. Data analysis was performed using the open-source R programming language software R version 4.2.3. **Results**

### Phenotypic resistance profile

Across the three ecological settings, *An. gambiae* (s.l.) populations displayed varying levels of resistance to the four tested insecticides (Fig 3A). Urban populations displayed incipient resistance to both organophosphates (pirimiphos-methyl) and carbamates (bendiocarb), with mortality rates of 96.9 ± 11.6% and 95.4 ± 13.2%, respectively. However, peri-urban and rural populations showed complete susceptibility to pirimiphos-methyl and bendiocarb, with mortality rates of 100%. Urban mosquitoes demonstrated slightly elevated resistance to deltamethrin (mortality rate: 40.7 ± 8.0%) compared to peri-urban and rural populations (mortality rates: 51.9 ± 9.7% and 55.4 ± 9.7%, respectively). Rural populations exhibited higher resistance to permethrin (mortality: 35.0 ± 4.0%) compared to urban (41.4 ± 6.9%) and peri-urban (43.7 ± 7.3%) populations.

**Figure 3.**
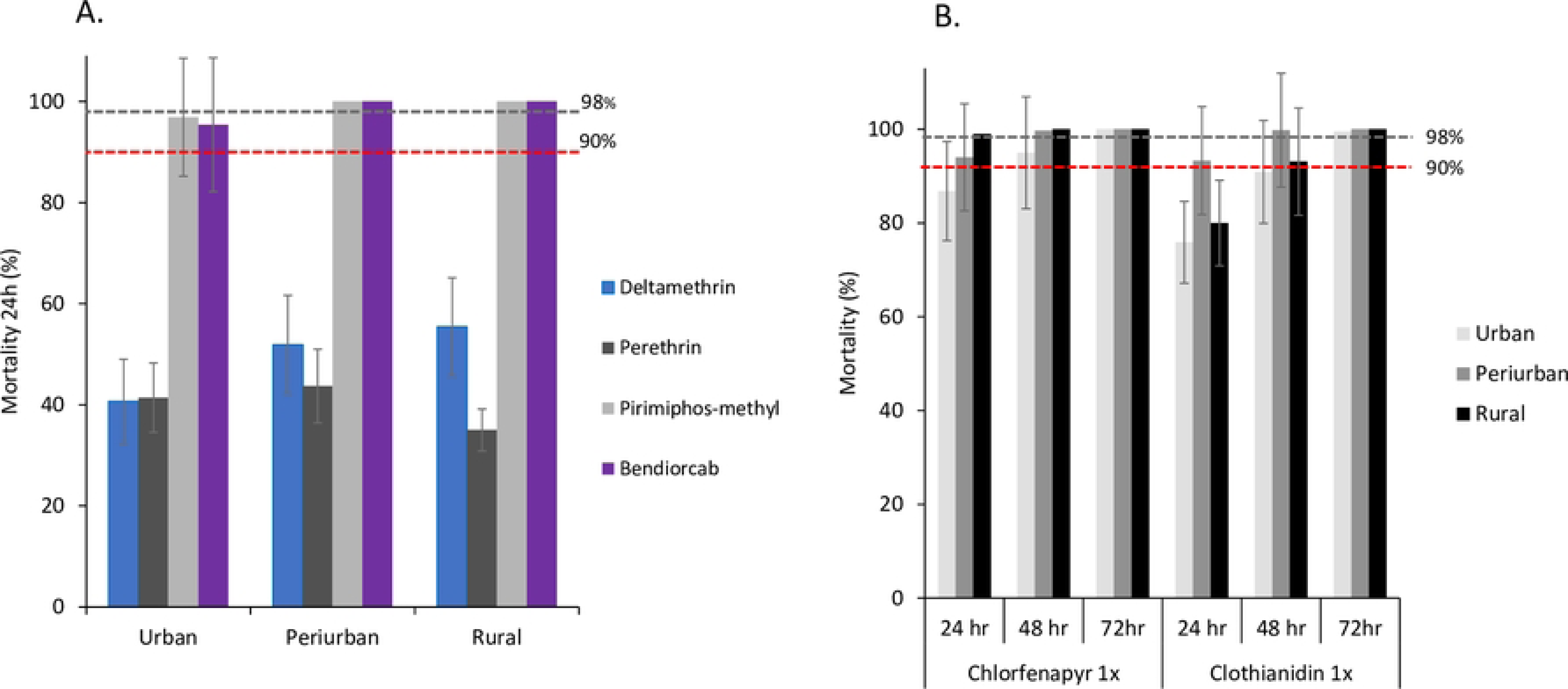
Mortality rates of *An. gambiae* s.l. from urban, peri-urban and rural areas in western Kenya. A) WHO tube bioassay; 24 hr. post-exposure to pyrethroids (deltamethrin and permethrin), organophosphate (pirimophos-methyl) and Carbamate (bendiocard); B) WHO bottle bioassay: 24-78 hr. post exposure to Chlorfenapyr 1X and Clothianidin 1X. Error bars indicate 95% confidence intervals. The 90% mortality threshold (red line) for declaring suspected resistance and 98% mortality threshold (black line) for calling full susceptibility based on the WHO criteria are indicated.

Among the field-collected mosquitoes, rural populations exhibited susceptibility (Mortality rate : 100%) to chlorfenapyr, within 24 hours however the peri-urban exhibited susceptibility after 48hrs, and the urban mosquitoes at 72hrs. A similar trend was observed for clothianidin, where mortality rates were below the susceptibility threshold (>98%) for all three populations within 48 hours after exposure. However, 100% mortality was observed within 72 hours (Fig 3B). The reference strain (Dongola) exhibited susceptibility to all insecticides, demonstrating 100% mortality at WHO-recommended discriminating dosages without significant variation, there by validating the quality of the test papers.

### Detection of detoxification enzymes with PBO synergist

Pre-exposure of *An. gambiae* (s.l.) populations to 4% PBO synergist underscored the involvement of monooxygenase enzyme activity in the pyrethroid detoxification, with mortality rates increasing to 88.5 ± 12.4% (urban), 90.3 ± 12.9% (peri-urban), and 87.6 ± 12.2% (rural) for PBO-deltamethrin. These results indicate a partial recovery of susceptibility by 47%, 40%, and 32%, respectively. Similarly, mortality rates increased to 78.8 ± 11.6% (urban), 90.7 ± 12.1%, and 84.2 ± 11.7% (rural) for PBO-permethrin, demonstrating partial recovery of susceptibility by 37%, 47%, and 49%, respectively. However, despite this partial recovery, the mortality rates after exposure to PBO synergist remained below the resistance threshold in all tested sites (Fig 4).

**Figure 4.**
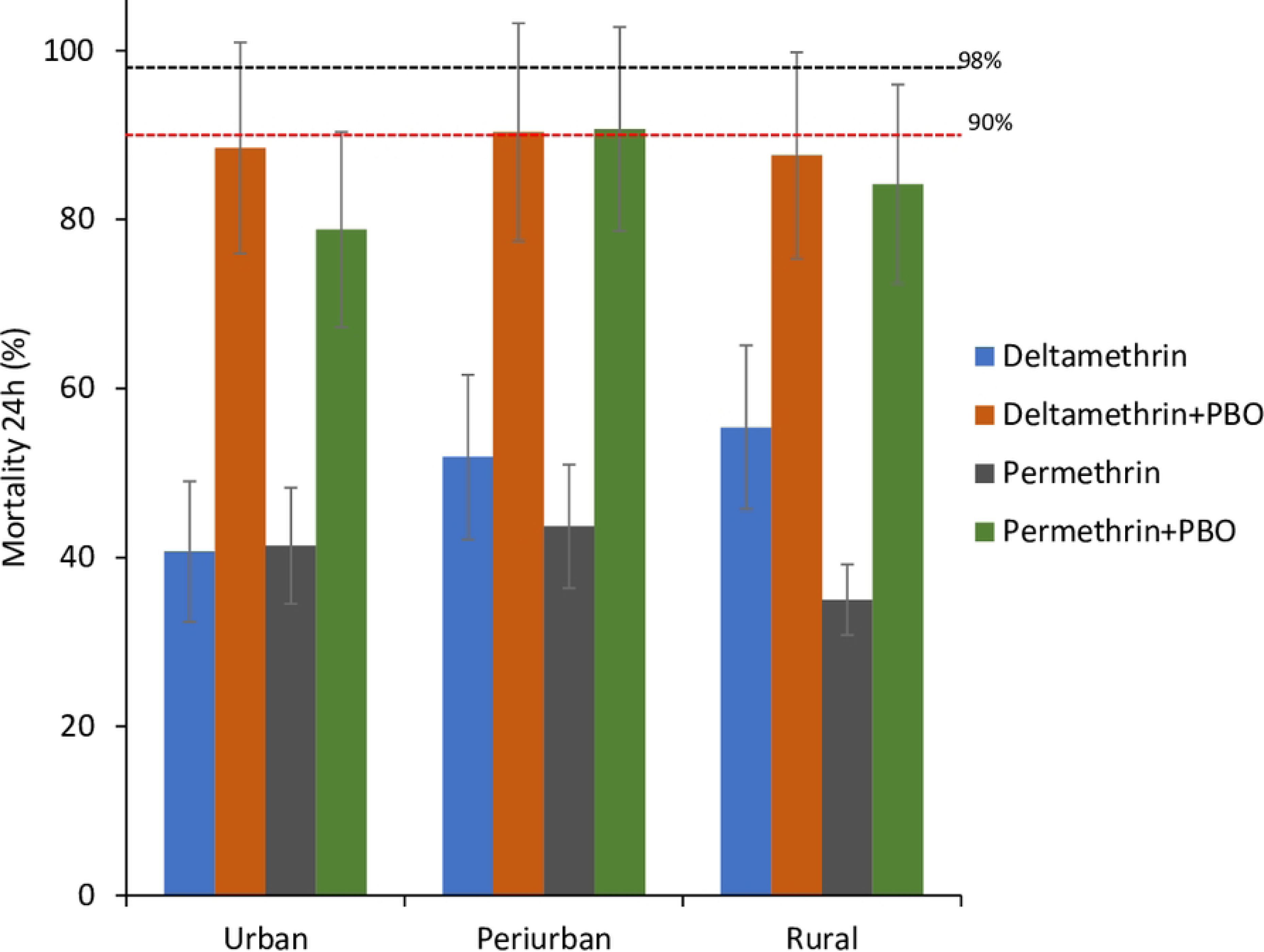
Synergist susceptibility profile of *Anopheles gambiae* (s.l.) populations from urban, peri-urban and rural zones after 60 min exposure to permethrin and deltamethrin alone and combined with PBO. Data shown as mean mortality with 95% confidence intervals. The 90% mortality threshold (red line) for declaring suspected resistance and 98% mortality threshold (black line) for calling full susceptibility based on the WHO criteria are indicated.

### Intensity of resistance

Following the baseline resistance against the pyrethroids, the escalation of deltamethrin resistance was tested at 5x and 10x concentrations. Urban *An. arabiensis* exhibited resistance to both concentrations, with mortality rates of 79.7 ± 13% at 5× and 85.2 ± 13.4% at 10×, indicating a high-intensity resistance (Fig 5). The mortality rates for peri-urban and rural populations exposed to 5× concentration was 93.8 ± 13.9% and 93.3 ± 13.4%, respectively. At 10× concentration, the mortality for both populations exceeded 98%, suggesting a moderate intensity of resistance.

**Figure 5.**
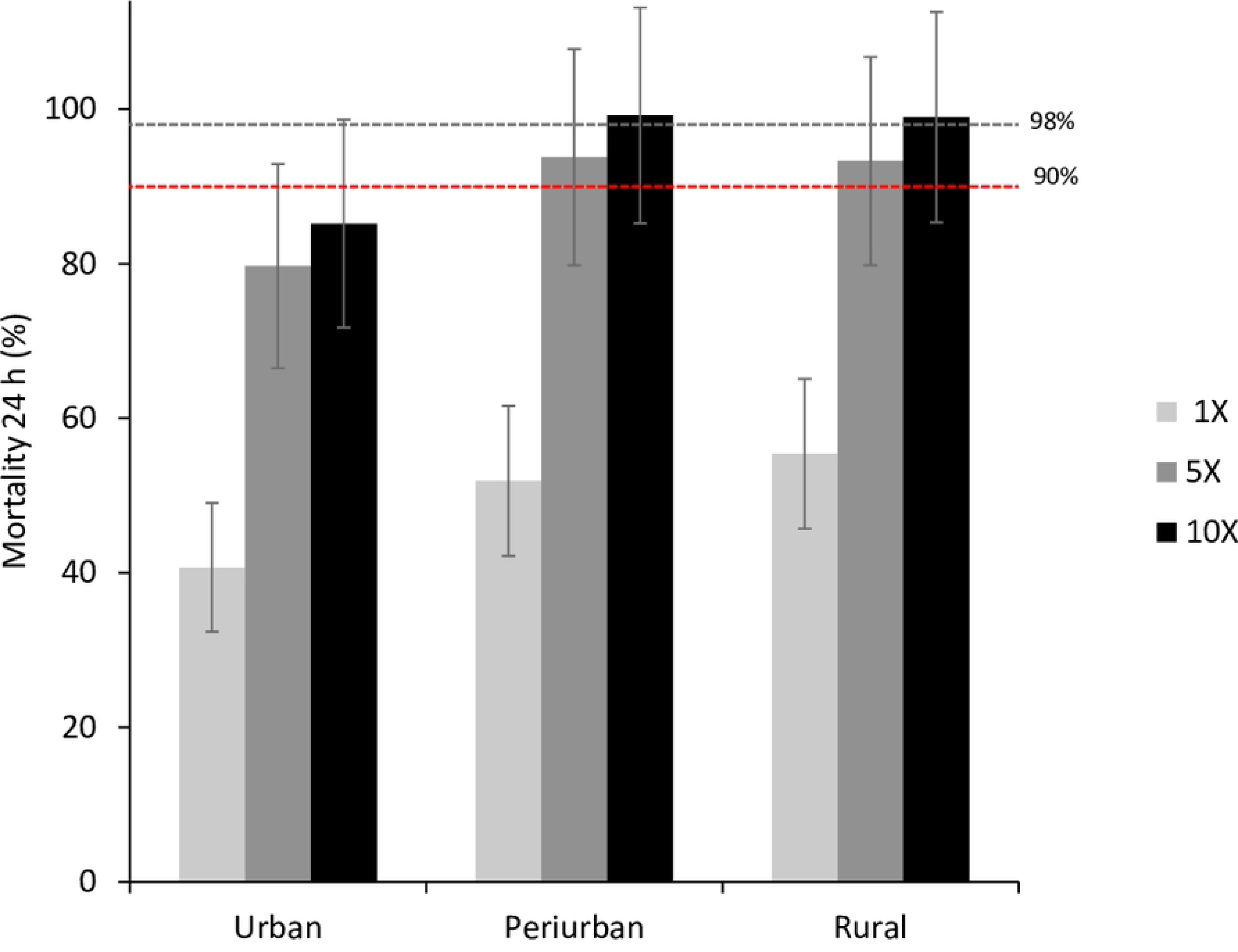
Insecticide resistance intensity status of *Anopheles gamabiae* (s.l.) from urban, peri-urban and rural areas. Standard resistance diagnostic concentration (0.05% deltamethrin, or 1X), 5X and 10X concentrations of deltamethrin were used WHO insecticide susceptibly tube bioassay. Error bars indicate 95% confidence intervals. The 90% mortality threshold for declaring suspected resistance and 98% mortality threshold () for calling full susceptibility based on the WHO criteria are indicated.

### Species composition

A sub-sample of randomly selected *An. gambiae* (s.l.) (1,907) underwent sibling species identification (urban 857, peri-urban 537 and rural 513). Within the *An. gambiae* (s.l.) samples, two species were identified: *An. gambiae* (s.s.) (henceforth *An. gambiae*) and *An. arabiensis*. *Anopheles arabiensis* was the predominant species in urban (96.3%, n=825) and peri-urban (96.8%, n=520) sites. Conversely, the rural populations were predominantly *An. gambiae* (82.7%, n=424) with few *An. arabiensis* (17%, n=89).

### Frequency of *Vgsc* and *Ace*-1 mutation

The L1014F mutation was detected at a high frequency in *An. arabiensis* specimens from urban sites, with an allelic frequency of 0.22, compared to peri-urban (0.11) and rural (0.14) populations. Hardy-Weinberg analysis revealed a significant deviation (P < 0.0001) from equilibrium in the urban population among the three *An. arabiensis* populations tested (Table 1). For *An. gambiae* samples tested from rural areas, the occurrence of the L1014F mutation was notably low at 0.01, while absent in peri-urban areas. Conversely, the L1014S mutation was detected in *An. gambiae*, with markedly high allelic frequencies in both rural (0.93) and fixed in peri-urban (although the number tested was very small, n=14). This mutation was also detected in *An. arabiensis* with very low allelic frequencies in urban (0.02) and peri-urban (0.01) areas, while in rural areas, it was present at a frequency of 0.21 (although the number tested was very small n=34). None of the An.

**Table 1.**
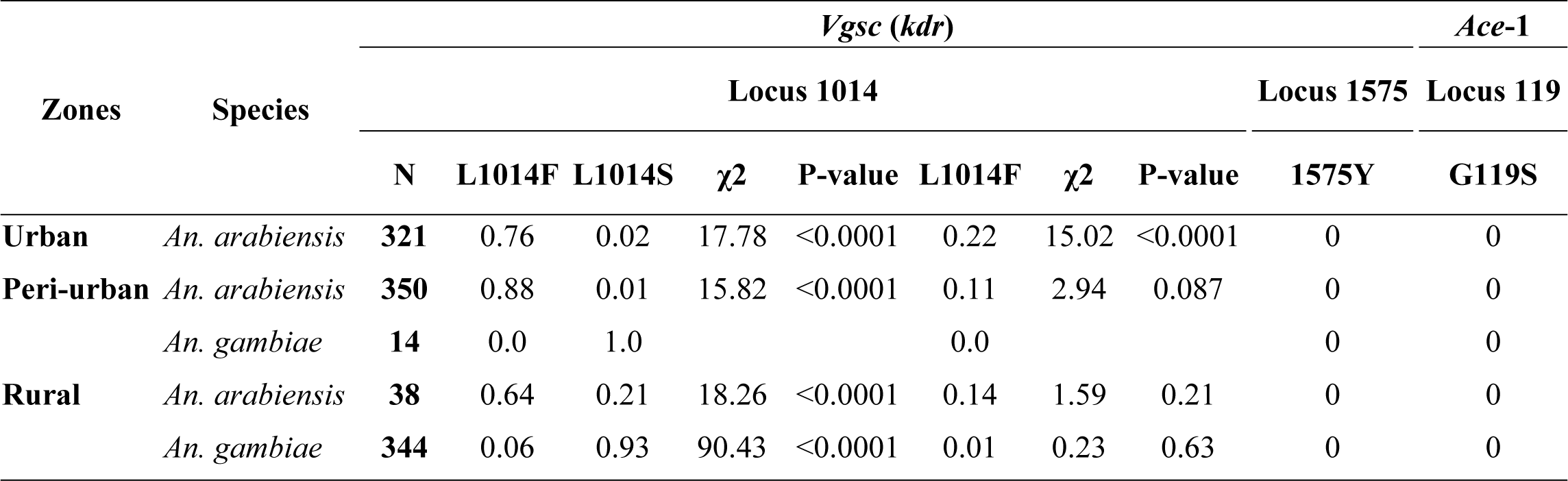
Allele frequency of *Vgsc (kdr)* and *Ace*-1 genes in the population of *An. arabiensis* and *An. gambiae* from urban, peri-urban, and rural zones in western Kenya.

*arabiensis* and *An. gambiae* populations tested for 1014S appeared to be in Hardy-Weinberg equilibrium (P < 0.001), possibly due to heterozygosity deficiency. The 1575Y of *Vgsc* and *Ace*-1 mutations were not detected in any of the populations tested.

### Detoxification enzyme activity

Biochemical assays revealed elevated enzymatic activities of mixed function oxidase (MFO) and non-specific esterases in field populations compared to the susceptible *Anopheles arabiensis* Dongola strain (Fig 6). Mixed function oxidase activity was notably higher in urban (mean activity = 0.8/mg protein), peri-urban (mean activity = 0.57/mg protein), and rural (mean activity = 0.6/mg protein) populations compared to the Dongola strain (0.43/mg protein) (χ2 = 99.775, *df* = 3, P < 0.0001; Fig 6A). Urban populations exhibited a 2-fold increase in MFO activity, while peri-urban and rural populations showed a 1.3-fold increase, all significantly higher than the Dongola strain (P < 0.0001). Pairwise comparisons indicated no significant difference in MFO activity between rural and peri-urban populations (Dunn’s test, P > 0.05).

**Figure 6.**
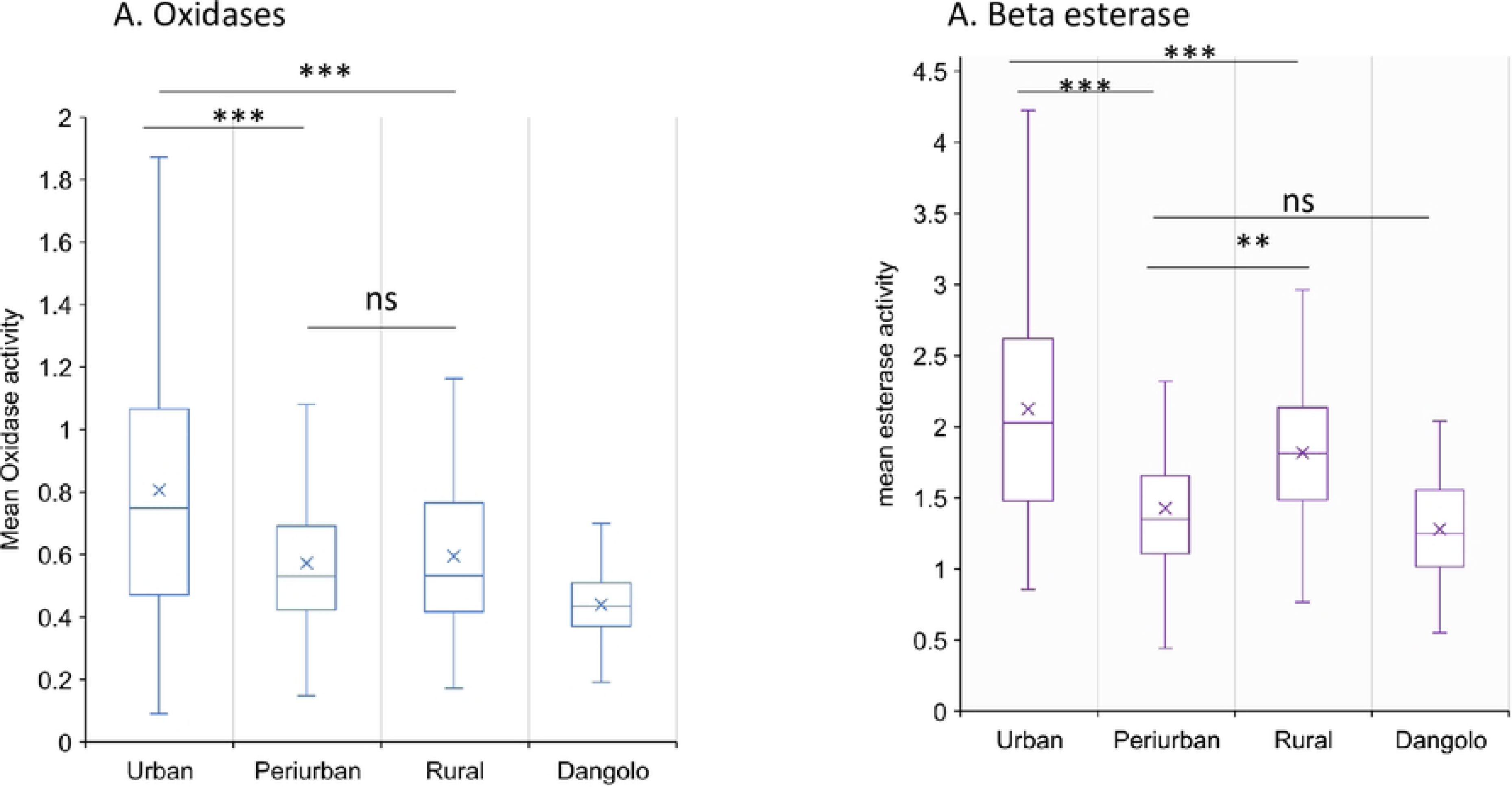
Metabolic detoxification enzymes activity in *Anopheles gambiae* (s.l.) in urban, peri-urban and rural zones A: monooxygenases; and B: β-esterase;. Error bars indicate 95% confidence intervals. *, P < 0.05; ***, *P* < 0.001; ns, not significant.

Notably, significantly increased esterase activity (using β-naphthyl acetate as a substrate) was observed in urban and rural populations compared to the Dongola strain (χ2 = 176.02, *df* = 2, P < 0.0001). Urban populations had a >1.2-fold increase in non-specific β-esterase activity (mean activity = 2.12/mg protein) compared to peri-urban (mean activity = 1.5/mg protein; Dunn’s test, p < 0.0001) and rural (mean activity = 1.82/mg protein; Dunn’s test, P < 0.0001) populations. However, no significant difference was detected in non-specific β-esterase activity between peri-urban (mean activity = 1.5/mg protein) and the Dongola strain (mean activity = 1.2/mg protein) (Dunn’s test, P = 0.78; Fig 6B).

### Pesticides used for the control of pests

Among the 40 different formulations reported by farmers for agricultural and public health (specifically, bedbug control) purposes, it was observed that pesticide formulations were more frequently utilized in urban environments (65%; 95% CI: 50-78%) compared to peri-urban (27.5%; 95% CI: 13.6-41.3%) and rural (47.5%; 95% CI: 32-63%) settings. However, the reported total may underestimate the actual number due to many farmers not recalling all the pesticide names they typically use. Pyrethroids emerged as the predominant class of pesticides (39% urban, 36.4% peri-urban, and 36% rural), followed by organophosphates (11.5% urban, 9% peri-urban, and 11% rural) and neonicotinoids (11.5% urban and 18.2% peri-urban) (Table 1). Among the pesticides mentioned, insecticides (40%; 95% Cl: 28-52%) and acaricides (40%; 95% Cl: 28-52%) were the most commonly reported in urban areas, whereas in peri-urban (53.5%; 95% CI: 42-63%) and rural (71.4%; 95% CI: 62-81%) settings, their usage was even more prevalent (Supplementary Table 1). Some respondents indicated using a combination of insecticides + acaricides (17.5%; 95% CI: 9-26%) in peri-urban areas and (27.4%; 95% CI: 18-37%) in rural settings. Furthermore, most respondents reported employing pesticides whenever they encountered pests on the farm any time of the season (77.8%; 95% CI: 68-88% urban, 88.7%; 95% CI: 82-96% peri-urban, and 69%; 95% CI: 59-79% rural).

## Discussion

This study, conducted across the urban-to-rural continuum of Kisumu city, enables us to report, for the first time, the susceptibility level of the primary malaria vector *An. arabiensis* in an urban environment in western Kenya to classes of insecticides used in vector control. Overall, relatively high resistance to pyrethroids was observed in urban populations of *An. arabiensis*, characterized by significantly elevated detoxification enzyme activities (MFO and β-esterases) and a high allelic frequency of 1014F mutations compared to peri-urban and rural populations. Incipient resistance to bendiocarb (carbamates) and pirimiphos-methyl (organophosphate) was detected in urban populations. Previous studies have documented widespread resistance to pyrethroids in *An. gambiae* (s.l) from rural and peri-urban areas in western Kenya (26–28, 34). The presence of high pyrethroid resistance in urban settings confirms the spatio-temporal spread of pyrethroid resistance in urban areas.

Species identification revealed distinct distributions: *An. arabiensis* was abundant in the drier urban and peri-urban settings, while *An. gambiae* dominated in the wetter rural areas. Previous studies in the lowlands of western Kenya around the Lake Victoria region have reported the dominance of *An. arabiensis* in the region (33, 42). *Anopheles arabiensis* demonstrates better survivorship under drier conditions in lowlands compared to *An. gambiae*, possibly contributing to its prevalence in urban and peri-urban lowlands (43, 44). Notably, this species has been found adapting in more polluted breeding sites associated with urbanization, unlike *An. gambiae*, which prefers small, temporary, sunlit, and clear freshwater pools, typical habitats found in rural areas (43, 45–47).

Until recently, *An. arabiensis* was considered more susceptible to insecticides in western Kenya region compared to sympatric populations of *An. gambiae* (34). However, our findings indicate this species displays high tolerance to increased doses of deltamethrin (pyrethroids) in urban settings, with moderate resistance observed in peri-urban and rural populations. Studies from the region confirm a rising trend of pyrethroid resistance in *An. arabiensis*, although not reaching the levels observed in the urban environment (28, 31, 48). The adaptability of *An. arabiensis* to urban environments and its high tolerance to insecticides is worrying. It is preference for outdoor resting and feeding behaviors (49–51) raises concerns on the effectiveness of LLINs as the sole malaria control strategy in urban areas. This underscores the necessity for integrated vector control strategies, including larval source management, to tackle both indoor and outdoor mosquito-biting behaviors and the increased resistance observed.

Biochemical analysis confirmed the overactivity of mixed-function oxidases and esterases in urban populations, along with a high frequency of the *kdr* allele 1014F, while this allele was less prevalent in peri-urban and rural areas. Conversely, the *kdr* 1014S mutation, fixed in *An. gambiae* in western Kenya, was found at low frequency in urban and peri-urban *An. arabiensis* but was more prevalent in rural populations of *An. arabiensis* and *An. gambiae* corroborating previous studies (28, 30, 34). The significant partial restoration of susceptibility after preexposure to PBO indicates that metabolic resistance may be driving the high resistance levels observed in the three populations, unlike kdr mutations. The observed incipient resistance to organophosphates and carbamates might be induced by the overexpression of detoxification genes rather than target site resistance, as the *Ace1* mutation was absent in most sequenced samples. Enzyme systems such as monooxygenase and esterase-mediated resistance have been reported to confer reduced susceptibility to malaria vectors against carbamates, organophosphates, and pyrethroids (52). Notably, all three populations remained fully susceptible to neonicotinoids (clothianidin) and pyrroles (chlorfenapyr), suggesting these classes hold promise for combating malaria vectors in the region despite the high levels of insecticide resistance observed.

Resistance to pyrethroids in the lowlands of western Kenya, including urban areas, may arise mainly from the extensive use of these insecticides in public health programs such as LLINs mass distribution campaigns. Deltamethrin and permethrin have long been utilized as primary active ingredients in LLINs across endemic regions in western Kenya (53). The use of these same compounds in agricultural practices, as evidenced in this study, could be driving the observed resistance in these populations. Additionally, the emergence of bed bug infestations has led to the use of new classes of insecticides for pest control in urban settings. This is concerning as it may promote resistance to the recently approved chemicals for vector control. Similar trends have been noted in most African countries(54–57). Although this study did not analyze the level of pollution in different breeding habitats in urban areas, it was evident that aquatic habitats where *An. gambiae* predominated in rural areas were clean-water pools, whereas urban breeding sites where *An. arabiensis* larvae were prevalent were polluted water pools and ditches contaminated with effluents from residential houses and roads. Adaptation to these polluted habitats may have contributed to the observed increase in detoxification enzymes in urban mosquito populations.

## Conclusion

Urban *Anopheles arabiensis* populations exhibit high pyrethroid resistance intensity, characterized by elevated detoxification enzyme activities (MFO and β-esterases) and 1014F mutations. Possible exposure to pesticides from agriculture and adaptation to urban pollution may have contributed to this resistance. Given the species’ unique behavior and ecology compared to *Anopheles gambiae*, tailored vector control strategies are needed to address this concern as the human population increase in urban settings.

## Declarations

### Ethics approval and consent to participate

The study was approved by the Maseno University Ethics Review Committee (MUERC Protocol No. 00456) and the University of California, Irvine Institutional Review Board (UCI IRB) and received authorization from the Ministry of Health, Kenya. Written informed consent was sought from household heads before data were collected from the households. All experiments and methods were carried out in accordance with the relevant guidelines and regulations of MUERC and UCI-IRB.

### Availability of data and materials

The dataset supporting the conclusions of this article is included within the article and its supplementary information files.

### Authors’ contribution

MGM, EO, YAA and GY conceived and designed the study. IN, MGM, SAO, BO participated in the field work and laboratory analysis, MGM did data analysis and drafted the manuscript. MCL determined the study site demarcations, MGM, HA, JG and CW supervised data collection and edited the draft manuscript writing. The final manuscript was edited by AG, EO, YAA and GY. All authors read and approved the final version of the manuscript.

## Acknowledgments

The authors wish to thank the volunteers for their participation in this study and the leadership of Kisumu County for allowing us to conduct the study in the area. We acknowledge the Entomology

Laboratory at Kenya Medical Research Institute, Kisumu, the field assistants in urban, peri-urban and rural areas for providing technical support.

## Abbreviations

*Vgsc*: Voltage gated sodium channel
*Ace*: angiotensin converting enzyme
*Kdr*: knock down resistance
PBO: piperonylbutoxide
PBO: piperonyl butoxide
MFO: mixed function oxidase
LLIN: long-lasting insecticidal nets
IRS: indoor residual spray

